# Computational Analysis of Single Nucleotide polymorphisms (SNPs) in Human TCell Acute Lymphocytic Leukemia Protein 1 (TAL1) Gene/Comprehensive Study

**DOI:** 10.1101/447540

**Authors:** Shaza W. Shantier, Hashim E. Elmansi, Mihad E. Elnnewery, Hind K. Osman, Isam-Aldin A. Alhassan, Fatima A. Abdelrhman, Ahmed A. Yagaub, Einas M. Yousif, Alaa I. Abdalla, Rawan A. Elamin, Howina S. Fadol, Afra A. Fadl Alla, Mohamed A. Hassan

## Abstract

**Background:** *TAL1* is a proto-oncogene whose distorted modifications in committed T-cell Precursors is related with the development of T-ALL, it also found to be related to many other human hematological diseases such as lymphoblastic lymphoma, immunodeficiency 18, acute myeloid leukemia and diamond-blackfan Anemia.

**Objectives:** This study aims to predict the effect of nsSNPs on *TAL1* protein structure function

**Methods:** Retrieved nSNPs in the coding and *3’UTR* regions were analyzed using different in silico tools. Interactions of *TAL1* with functionally similar genes were investigated using Genemania. Post-translational modifications in several sites of the protein were also investigated.

**Results:** Out of ninety nsSNPs identified, only eight were found damaging to protein function of which one is located in the basis helix-loop-helix domain (bHLH). Two SNPs were anticipated by PolymiRTs to prompt disturbance or creation of miR binding sites.

**Conclusion:** The present study is the first ever computational analysis of *TAL1*’s nsSNPs hence this effort might be of help in the near future for inventing early diagnostic and therapeutic measures for T-ALL

## Introduction

Acute lymphoblastic leukemia (ALL) is a malignant disease of the bone marrow in which hematopoietic stem cells (HSC), which are called lymphoid cells in this case, are arrested and transformed in an early stage [1,2]. ALL accounts for 26% of childhood malignancy, making it the most common cancer in that age group [3]. There are two subtypes of the disease T-cell ALL (T-ALL) and B-cell precursor ALL (BCP-ALL), T-ALL involves thymocytes and responsible for about 15% of childhood and 25% of adult ALL [4]. *TAL1* was also found to be related to many human hematological diseases such as lymphoblastic lymphoma, immunodeficiency 18, acute myeloid leukemia and diamond-black fan Anemia [5-9].

Although the local alterations or chromosome translocation in committed T-cell precursors is associated with the development of T-ALL [10-12], literature review revealed that there are no previous studies on the effect of such alterations on *TAL1* structure and function. Therefore the aim of the present work was to conduct a full in silico analysis of *TAL1*’s SNPs using bioinformatics prediction tools to study the possible effect of the genetic variations on the protein structure and function.

Studies showed that tumor-specific activation of *TAL1* is found in 40-60% of T-ALL patients, resulting from interstitial chromosome mutations (25-30%), chromosomal translocation (4-5%) or by undefined mechanisms (60%) [13-16]. These mutations are expected to be because of single nucleotide polymorphisms (SNPs), which are the most well known kind of genetic variation among individuals, consisting about 90% of genetic polymorphisms [17]. One type of SNPs is the non-synonymous SNPs (nsSNPs), also known as missense SNPs, which are very important because they are responsible for changes in human proteins’ functions by substituting amino acid residues [18].

Therefore, early prediction and better understanding of the *TAL1* gene functions could help in improving the prognosis of the disease. As this analysis is the first precise and broad computational study of functional SNPs in the *TAL1* gene, it might be of great help in the near future for inventing early diagnostic and therapeutic measures for T-ALL.

## Materials and Methods

### SNPs Retrieval

Polymorphisms in *TAL1* gene were retrieved from the national center of biotechnology information database; dbSNP/NCBI database NCBI [19]. The retrieved SNPs were then filtered for investigation.

### Insilico analysis of non-synonymous single nucleotide polymorphisms (nSNPs)

Different soft-wares were utilized to study the impact of SNPs mutations on *TAL1* protein structure and function. Deleterious effect of nSNPs was investigated by SIFT and Polyphen-2 softwares. Stability changes was explored by I mutant-3. The association of nsSNPs with disease was done by PhD-SNP programming. The auxiliary changes in 3D structure were dissected utilizing Chimera programming. SNPs at the 3’UTR were likewise broke down to identify the impact on microRNA restricting destinations utilizing PolymiRTs programming. *TAL1* gene interactions were investigated using GENEMANIA. In this study, nsSNPs and those at the *3’UTR* regions were selected for analysis.

### Investigation of *TAL1* Gene’s Interactions and Appearance in Networks in GENEMANIA Database

The online database GENEMANIA studies the gene function and interrelation with other genes using functional association data including protein and genetic interactions, pathways, co-expression, colocalization and protein domain similarity. It can also be used to find new members of a pathway or complex, find additional genes that may have been missed in screening or find new genes with a specific function, such as protein kinases [20]. Available at: http://www.genemania.org/.

### nsSNPs’ Structural impact

Functional effects of nsSNPs were analyzed using SIFT (http://sift.bii.a-star.edu.sg/) in which SNPs are characterized into tolerated and deleterious. The input SNPs’ rs-IDs were submitted to the server for analysis, prediction was given as a tolerance index (TI) score going from 0.0 to 1.0. SNPs with TI score under 0.05 were anticipated to be deleterious; those more prominent than or equal to 0.05 were anticipated to be tolerated (http://blocks.fhcrc.org/sift/SIFT.html) [21].

### Prediction of Deleterious nsSNPs

Polyphen software (Polymorphism Phenotyping v2; http://genetics.bwh.harvard.edu/pph2) calculates position-specific independent count (PSIC) scores of which 1.0 is considered to be damaging. The SNPs are appraised quantitatively as benign, possibly damaging and probably damaging [22]. Positions of interest and new residue in protein FASTA sequence were submitted to Polyphen to investigate the functional effect of mutations.

Damaging SNPs in the above softwares were further analyzed by I mutant and PhD servers to estimate their effects on protein stability and disease associated variations, respectively.

### nsSNPs’Impact on Protein Stability

#### I-Mutant3.0 server

I-Mutant is a web server for the automatic prediction of protein stability changes upon single-site mutations starting from the protein structure or sequence. It calculates the free energy change value (DDG) and predicts the indication of the free energy change value (DDG) (increase or decrease), along with a reliability index for the results (RI: 0–10, where 0 is the minimum reliability and 10 is the maximum reliability). A DDG < 0 corresponds to a decline in protein stability, whereas a DDG > 0 corresponds to an increase in protein stability [23]. The residues changes and protein sequence in FASTA format were submitted to I-mutant server to process DDG value (kcal/mol) and the RI value. Conditions for all enteries were set at temperature 25°C and pH 7.0. Available at http://gpcr2.biocomp.unibo.it/cgi/predictors/I-Mutant3.0/IMutant3.0.cgi.

### Prediction of Disease Associated Variations

PhD-SNP is Support Vector Machine based classifier that predicts the disease associated variations upon single point mutation [24-26]. The residues changes and protein sequence in FASTA format were submitted to PhD-SNP server for the analysis. Available at: http://snps.biofold.org/phd-snp/phdsnp.html.

### Prediction of the Impact of SNPs at the 3Un Translated Region (*3’UTR*) by PolymiRTS Database

PolymiRTS (v3.0) database is designed specifically for the analysis of non-coding SNPs at *3’UTR* and identification of nSNPS that affect miRNA (micro RNA) targets in human and mouse [27]. *3’UTR* SNPs in *TAL1* gene were analyzed in order to investigate the alteration of miRNA binding on target sites which may result in diverse functional consequences. Available at: http://compbio.uthsc.edu/miRSNP.

### Project hope

HOPE is an online service, developed at the Centre for Molecular and BiomolecularInformatics CMBI at Radboud University in Nijmegen. It gathers structural information from different sources, including calculations on the 3D protein structure, succession comments in UniProt and expectation from the Reprof software. HOPE consolidates this data to analyze the impact of specific mutation on the protein structure [28].

### Homology modeling

The 3D models for protein wild type and mutated were produced utilizing two homology modeling portals; Phyre2 and Raptorx [29, 30]. The obtained structures were then visualized by Chimera 1.10.2

### UCSF Chimera

It is an extensible program for interactive representation and investigation of molecular structures and related information, including density maps, supramolecular gathering, sequence arrangement, docking results and conformational ensembles [31]. Chimera [32] can give the 3D structure of the protein and then changing between wild and mutant amino acids with the candidate to show the resulted effect. Chimera accepts the input in the form of pdb ID or pdb file. (https://www.cgl.ucsf.edu/chimera/).

### Predicting post translational modification (PTM) sites

The phosphorylation sites of *TAL 1* at serine, threonine and tyrosine residues were predicted by NetPhos server [33]. The ubiquitylation sites at lysine residue were investigated by UbPred and BDM-PUB tools [34].

## Results

## Discussion

Genetic polymorphism in *TAL1* gene was found sharing protein domain (bHLH domain) with five genes and associated in many diseases such as lymphoblastic leukemia, craniosynostosis, anemia and fibrodysplasia ossificans progressive.

Our present study detected eight SNPs in *TAL1* coding region to be highly damaging. The interesting finding was that the clinical significance for the four disease-related SNPs was unknown in the dbSNP/NCBI database. Insilico analysis of single nucleotide polymorphisms (SNPs) has become a valuable and essential tool for prediction of variants most likely associated with disease. This approach has been done for many disorders especially for cancer related genes [35-38]. Our methods in bioinformatics analysis were in conjunction with previous papers analyzing disease-related genes such as *VCAM-1*, *MSH6*, *GRM4*, *MYC* and *TAGAP* genes [39-43].

*TAL1* has similar expression to four genes which are mainly either inhibitors of protein binding or regulating transcription binding factors (Figure 1, Table 1). It has many molecular functions; it enables RNA polymerase II transcription factor activity and colocalizes with histone deacetylase complex. HSCs undergo differentiation when TAL1 activates transcription by recruiting a core complex DNA consisting of *E2A*/*HEB*, *GATA1/2/3*, *LMO1/2*, *LDB1*, and an additional complex comprising *ETO2*, *RUNX1*, *ERG* or *FLI1* [44]. The *TAL1*’s regulatory functions (lineage priming, activation, and repression of gene expression programs) give knowledge into principal developmental and transcriptional components and features mechanistic parallels between typical and oncogenic processes [45]. Also, Bae et al found that one intronic SNP of *TAL1* gene (rs2250380) was significantly associated with Schizophrenia [46].

**Figure 1:**
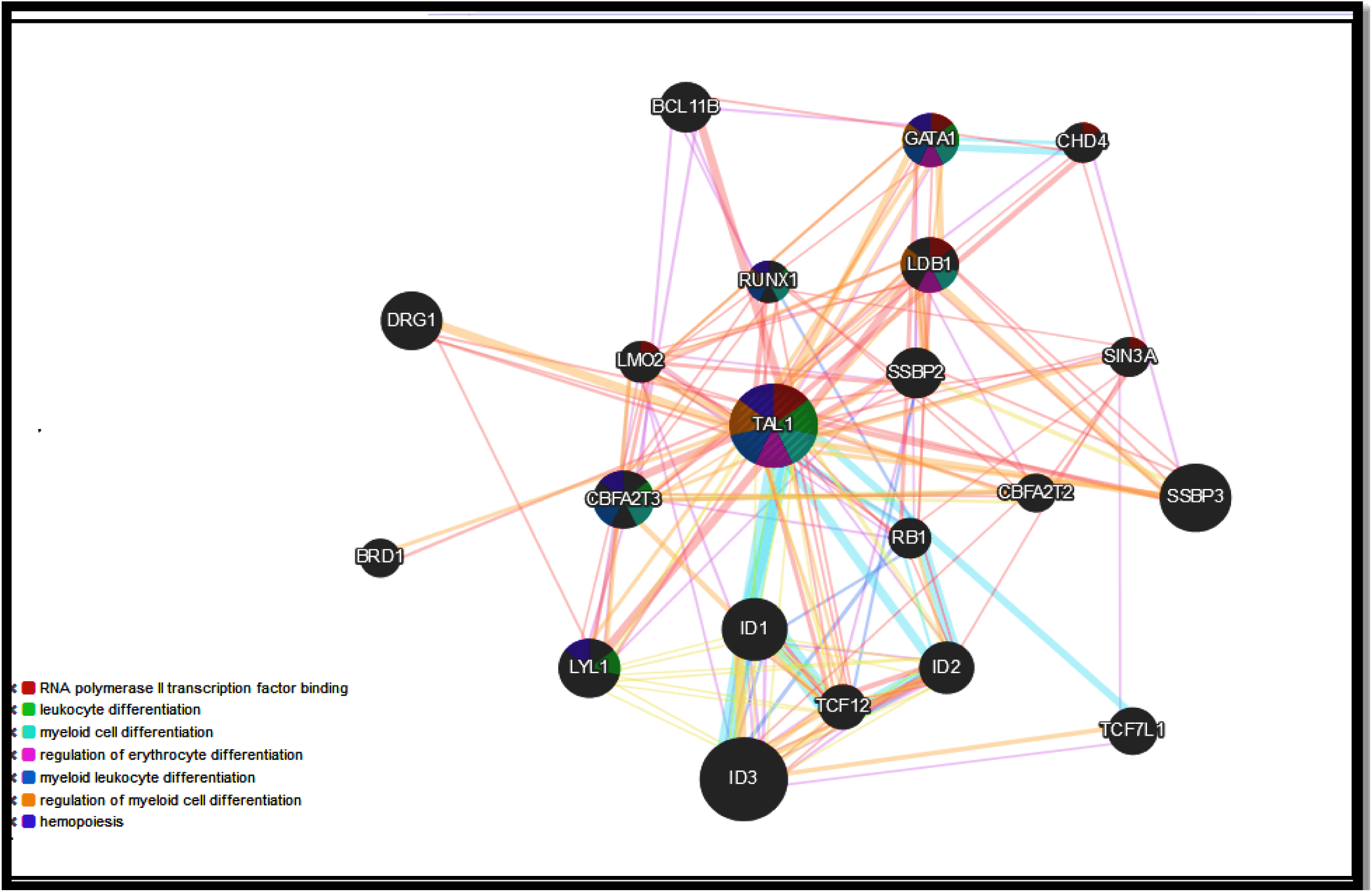
shows functional interaction between *TAL1* and its related genes.

**Table1.** Shows the genes co-expressed, physical interaction and share a domain with *TAL1*

| Gene symbol | Description | Co-expression | Physical interaction | shared domain |
| --- | --- | --- | --- | --- |
| *ID3* | inhibitor of DNA binding 3, HLH protein | NO | NO | YES |
| *SSBP3* | single stranded DNA binding protein 3 | NO | YES | NO |
| *ID1* | inhibitor of DNA binding 1, HLH protein | NO | NO | YES |
| *LYL1* | lymphoblastic leukemia associated hematopoiesis regulator 1 | NO | YES | YES |
| *DRG1* | developmentally regulated GTP binding protein 1 | NO | YES | NO |
| *CBFA2T3* | CBFA2/RUNX1 translocation partner 3 | NO | YES | NO |
| *LDB1* | LIM domain binding 1 | NO | YES | NO |
| *GATA1* | GATA binding protein 1 | YES | YES | NO |
| *ID2* | inhibitor of DNA binding 2, HLH protein | YES | NO | YES |
| *SSBP2* | single stranded DNA binding protein 2 | NO | YES | NO |
| *BCL11B* | B-cell CLL/lymphoma 11B | NO | YES | NO |
| *TCF7L1* | transcription factor 7 like 1 | NO | NO | NO |
| *TCF12* | transcription factor 12 | NO | YES | YES |
| *RUNX1* | runt related transcription factor 1 | NO | YES | NO |
| *LMO2* | LIM domain only 2 | YES | YES | NO |
| *RB1* | RB transcriptional corepressor 1 | NO | YES | NO |
| *CHD4* | chromodomain helicase DNA binding protein 4 | NO | YES | NO |
| *SIN3A* | SIN3 transcription regulator family member A | YES | YES | NO |
| *BRD1* | bromodomain containing 1 | NO | Yes | NO |
| *CBFA2T2* | CBFA2/RUNX1 translocation partner 2 | NO | Yes | NO |

**Table2.**
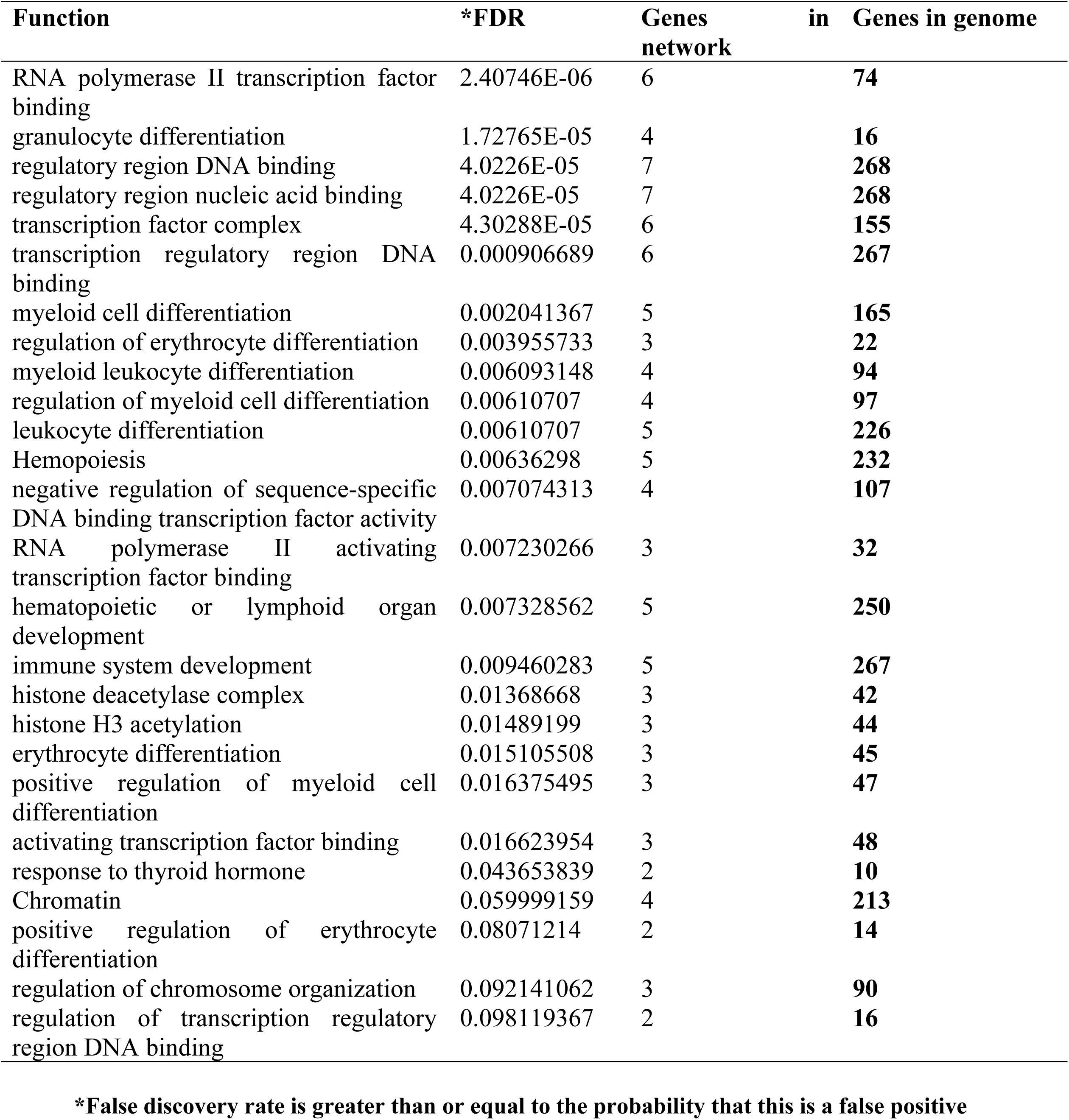
Illustrates the *TAL1* functions and its appearance in network and genome

Four thousand two hundred and twelve SNPs in *TAL1* gene were retrieved from dbSNP/NCBI in May 2018. There were nsSNPs, *3’UTR*, *5’UTR* and others (Figure 2). All nsSNPs in the coding region were subjected to different tools to test their effect on protein function and stability. Ninety nsSNPs were determined by SIFT to be tolerated or deleterious. According to Polyphen2, they were found to be benign, possibly or probably damaging. SNPs which scored ≤ 0.05 in SIFT and ∼1 in Polyphen-2 were then selected so that only the deleterious and damaging ones would be analyzed. The eight shortlisted damaging SNPs were further analyzed by I-Mutant to predict the effect of the mutant amino acids on the protein steadness. The obtained results reflected that the stability with related free energy will be different due to mutation. Six SNPs (M →R, F→ V, A→ T in different positions) had decrease effect on stability with DDG values ranged between −0.79 ‒ −1.08, while the two SNPs (S → Y; rs369207599) increase the stability of protein and also analyzed to be disease related using PHD- SNPs software (Table 3).

**Figure 2:**
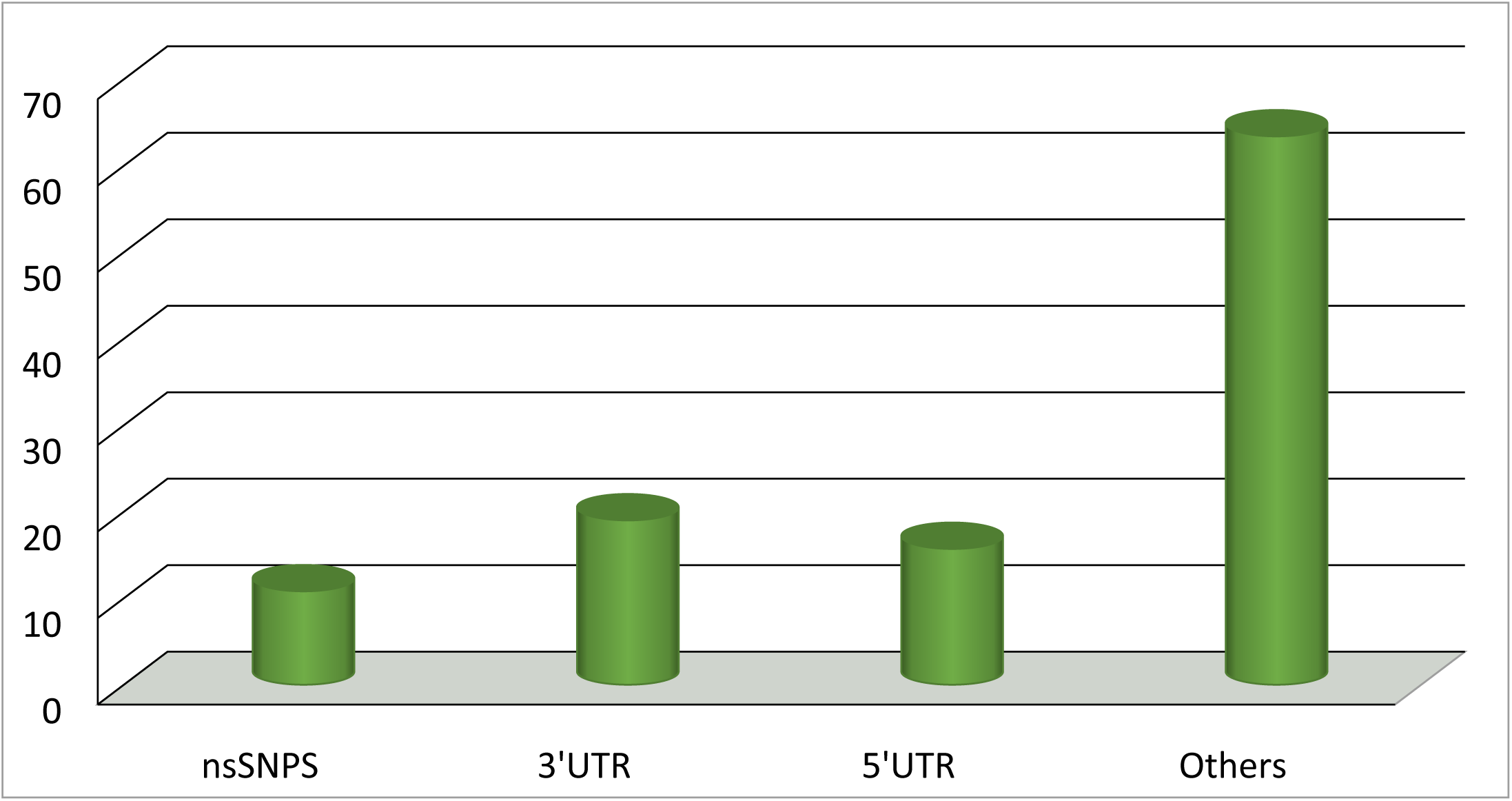
Percentages of the SNPs in *TAL1* gene. (nsSNPs: 10.8%; *3’UTR* SNPs: 19%; *5’UTR* SNPs: 15.7%; Other SNPs: 63%)

**Table 3.** Represents prediction results of SIFT, Polyphen, I mutant and PhD-SNP

| SNP | Amino Acid Change | Sift Score | Sift Prediction | Polyphen score | Polyphen prediction | I mutant |  | PhD-SNP |  |  |
| --- | --- | --- | --- | --- | --- | --- | --- | --- | --- | --- |
|  |  |  |  |  |  | SVM2 | DDG | R I | Effect | R I |
| rs147015002 | M160R | 0.009 | Deleterious | 0.835 | Possibly damaging | Decrease | -1.08 | 2 | Neutral | 1 |
| rs147015002 | M162R | 0.01 | Deleterious | 0.835 | Possibly damaging | Decrease | -1.08 | 2 | Neutral | 1 |
| rs200970768 | F151V | 0.013 | Deleterious | 0.615 | Possibly damaging | Decrease | -0.99 | 6 | Neutral | 5 |
| rs200970768 | F153V | 0.035 | Deleterious | 0.745 | Possibly damaging | Decrease | -1.07 | 6 | Neutral | 5 |
| rs369207599 | S172Y | 0.004 | Deleterious | 0.981 | Probably damaging | Increase | -0.29 | 2 | Disease | 1 |
| rs369207599 | S174Y | 0.006 | Deleterious | 0.939 | Probably damaging | Increase | -0.29 | 2 | Disease | 1 |
| rs375935172 | A210T | 0.006 | Deleterious | 0.97 | Probably damaging | Decrease | -0.79 | 8 | Disease | 5 |
| rs375935172 | A208T | 0.006 | Deleterious | 0.97 | Probably damaging | Decrease | -0.79 | 8 | Disease | 4 |

**\*DDG: free energy change value; SVM2: support vector machine; RI: reliability index**

The rs375935172 caused amino acid substitution A208T and A210T. The mutant residue is bigger and more hydrophilic than the wild-type residue, which may cause loss of interactions with other molecules on the surface of the protein. This mutation is the most important as it is situated in the bHLH domain. The change presents an amino acid with various properties, which can disturb this domain and abrogate its function. The mutated residue is situated in a domain that is essential for binding of other molecules. It is possible that the mutation affects these contacts and might influence the interaction disturb signal transfer from binding domain to the activity domain.

The rs147015002 results in the substitution of amino acid M to R at positions 160 and 162. The original wild-type residue and newly introduced mutant residue differ in their specific size, charge, and hydrophobicity-value. The mutant residue is bigger than the wild-type residue which might lead to bumps. The wild-type residue charge was neutral; the mutant residue charge is positive which can cause repulsion of ligands or other residues with the same charge. The wild-type residue is more hydrophobic than the mutant residue. Hydrophobic interactions, either in the core of the protein or on the surface, will be lost.

The rs200970768 caused conversion of amino acid F to smaller amino acid V at position 151 or 153, while the rs369207599 result in replacement of S into Y at position 172 or 174 which differ in size. The mutant residue is bigger and this might lead to bumps. The mutant residue prefers to be in another secondary structure; therefore, the local conformation will be slightly destabilized (Figure 3). Functional SNPs in three translated region in *TAL1* gene was analyzed using PolymiRTS software. Among 227 SNPs in *3’UTR* there were only 48 functional SNPs predicted (S1 Table). rs181722922 SNP contain (C) allele has 12 miRNA Sites as Target binding site can create a new microRNA site. rs145888818 SNP contain (D) allele has 10 miRNA Sites which they are derived allele that disrupts a conserved miRNA sit (Table 4).

**Figure 3.**
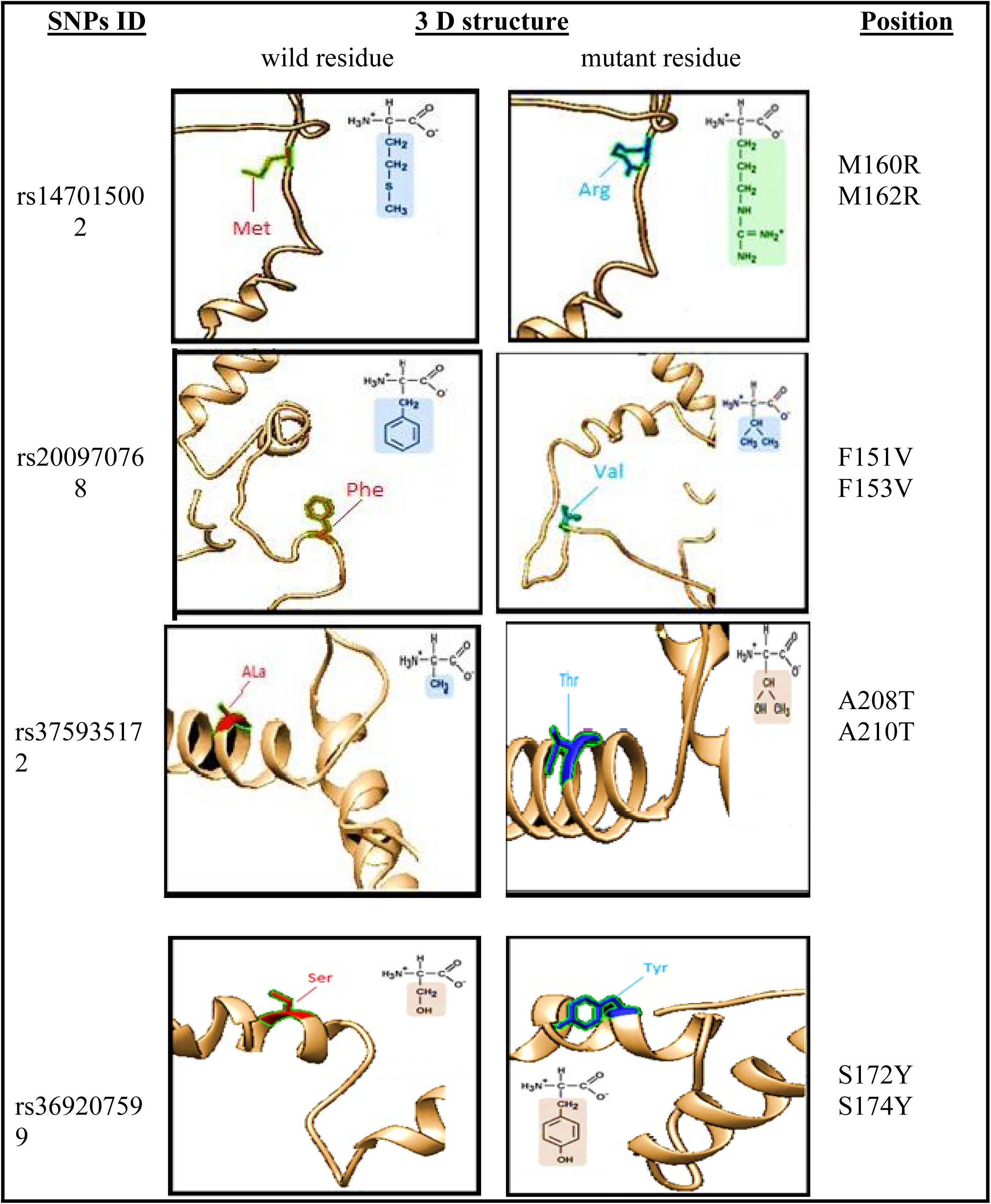
3D model of *TAL1* protein (visualized by Chimera 1.10.2).

**Table 4.**
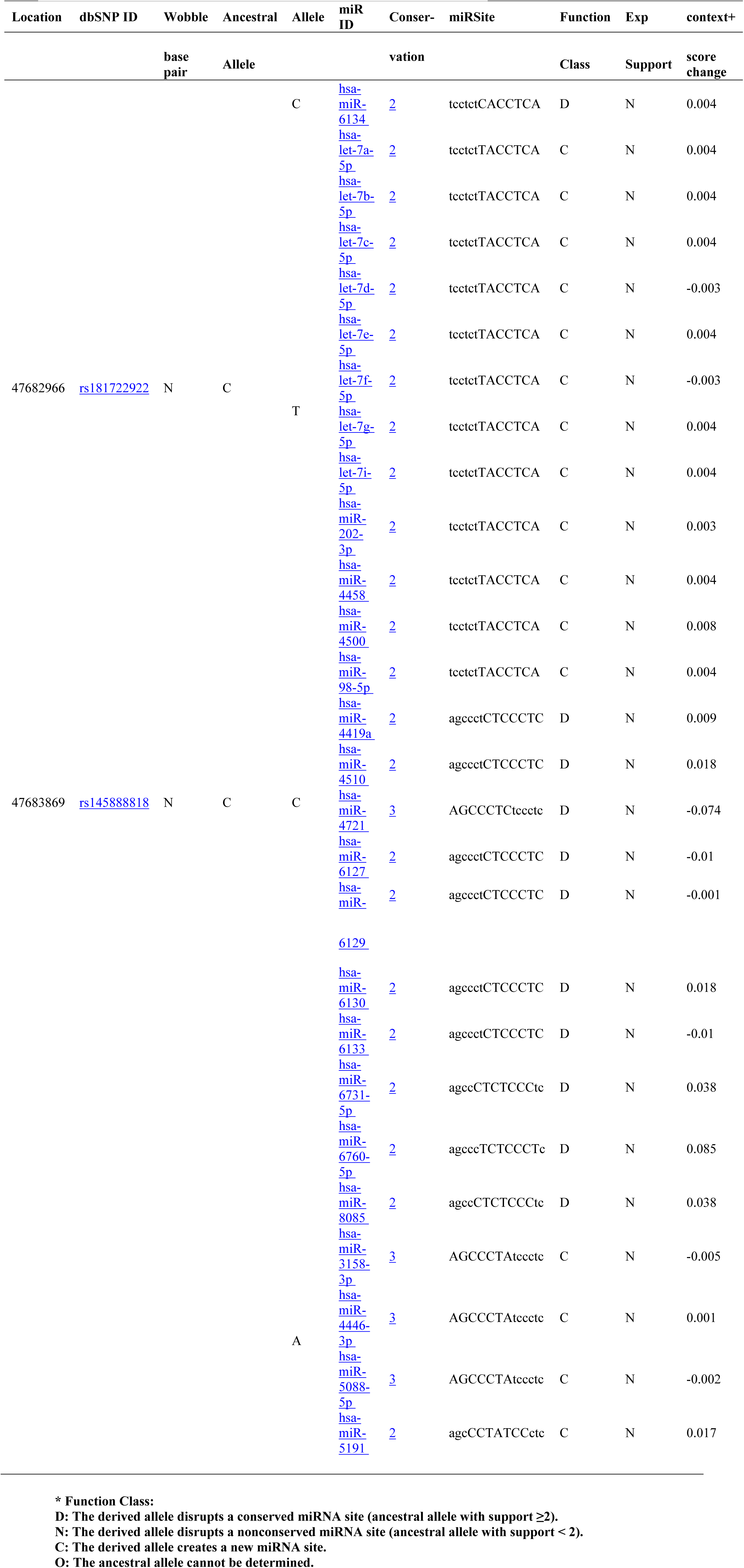
Shows SNPs and INDELs in miRNA target sites

PTMs are imperative in directing structures and functions of proteins hence are engaged in numerous biological events, for example, protein-protein interactions and cell signaling etc [47, 48]. Phosphorylation of proteins is an essential regulatory mechanism as it changes the structural conformation of a protein, resulting in making it to become activated, deactivated, or altering its function [49]. Target amino acid is usually serine, threonine or tyrosine residues. Net Phos predicted 15 Serine, 7 Threonine and 2 Tyrosine residues which have high potentiality to be phosphorylated (Table 5). Ubiquitination (or ubiquitylation) is an enzymatic post-translational modification in which an ubiquitin protein is linked to a substrate protein. It changes cellular process by regulating the decomposition of proteins (via the proteasome and lysosome), arranging the cellular localization of proteins, activating and inactivating proteins, and modulating protein-protein interactions [50-52]. UbPred and BDM-PUB tools predicted five and seven Lysine residues, respectively which undergo ubiquitylation (Table 6). Although PTMs are not coincided in position with the nsSNPs in *TAL1* gene, results by similarity revealed that phosphorylation of Ser-122 in *TAL1*gene is strongly stimulated by hypoxia and subsequently ubiquitination targets the protein for rapid degradation via the ubiquitin system (Table 6). This process may be characteristic for microvascular endothelial cells, since it could not be observed in large vessel endothelial cells.

**Table 5:**
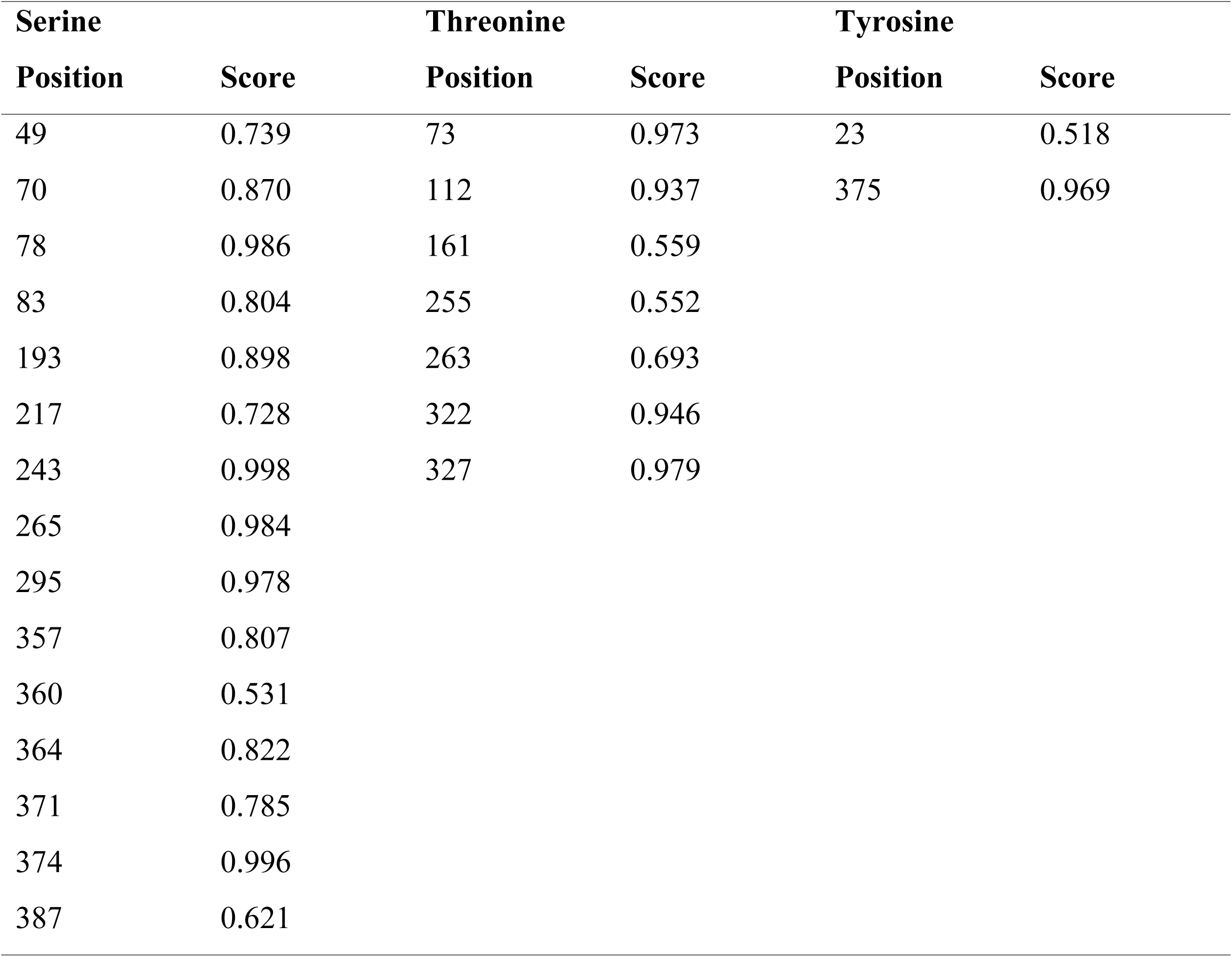
Phosphorylation sites predicted by NetPhos 3.1

| **Serine** |  | **Threonine** |  | **Tyrosine** |  |
| --- | --- | --- | --- | --- | --- |
| **Position** | **Score** | **Position** | **Score** | **Position** | **Score** |
| 49 | 0.739 | 73 | 0.973 | 23 | 0.518 |
| 70 | 0.870 | 112 | 0.937 | 375 | 0.969 |
| 78 | 0.986 | 161 | 0.559 |  |  |
| 83 | 0.804 | 255 | 0.552 |  |  |
| 193 | 0.898 | 263 | 0.693 |  |  |
| 217 | 0.728 | 322 | 0.946 |  |  |
| 243 | 0.998 | 327 | 0.979 |  |  |
| 265 | 0.984 |  |  |  |  |
| 295 | 0.978 |  |  |  |  |
| 357 | 0.807 |  |  |  |  |
| 360 | 0.531 |  |  |  |  |
| 364 | 0.822 |  |  |  |  |
| 371 | 0.785 |  |  |  |  |
| 374 | 0.996 |  |  |  |  |
| 387 | 0.621 |  |  |  |  |

**Table 6:**
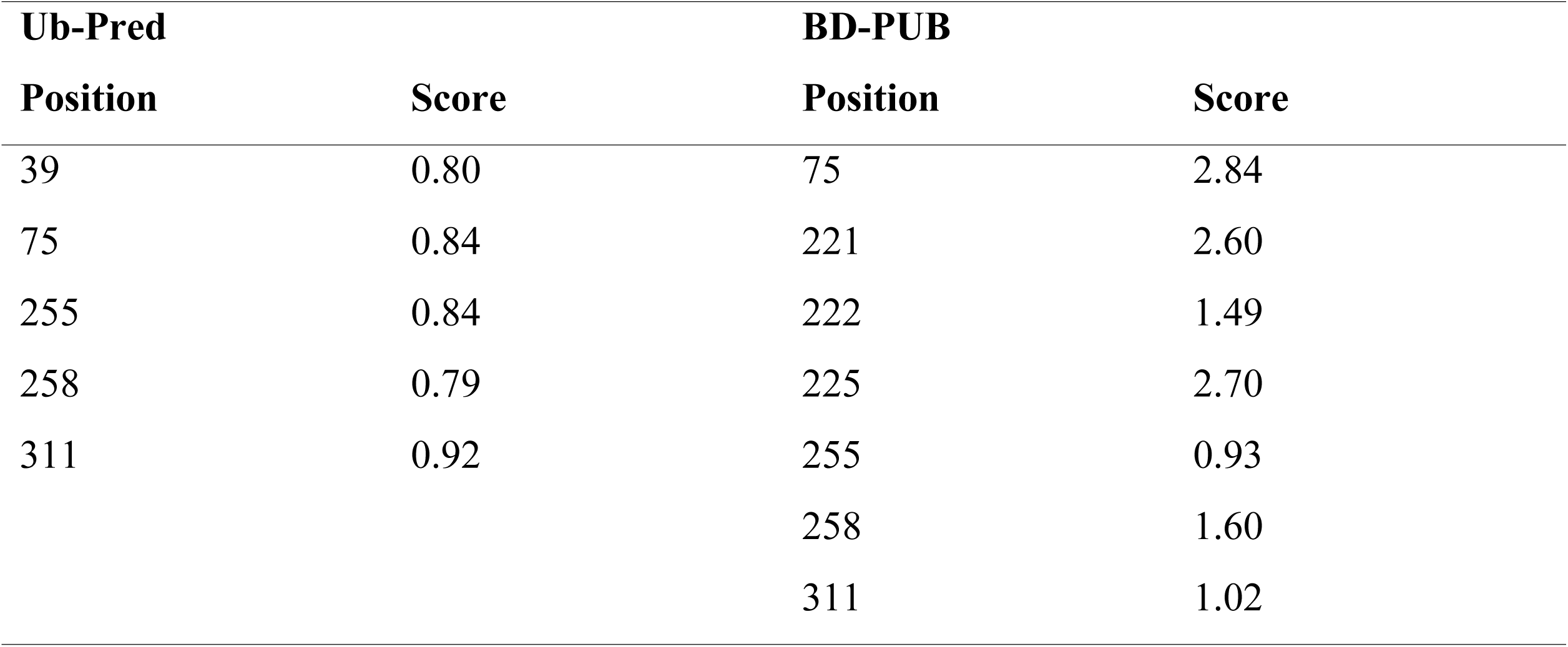
Ubiquitylation sites predicted by Ub-Pred and BDM-PUB

## Conclusion

This study predicted that the stability and function of *TAL1* protein is affected by eight high risk nsSNP. One of these mutationS (rs375935172) is located at the highly conserved bHLH domain region, hence it is of high concern as this is the only functional region of the protein. In addition to these findings, the study identifies several *TAL1* sites that undergo post transitional modification. Therefore, further investigations and wet lab experimentation are required to determine the effects of these polymorphisms on the protein function which can be a hit for discovering new drugs.

## References

1. ((AML): Practice Essentials, Pathophysiology, Etiology. [online] Emedicine.medscape.com. Available at: http://emedicine.medscape.com/article/197802-overview [Accessed 21 Apr. 2018].

2. Disease-ontology. Acute leukemia. [online] Available at: http://disease-ontology.org/term/DOID%3A12603/ [Accessed 8 May 2018].

3. Harisson CJ. Acute Lymphoblastic Leukemia. Clinics in Laboratory Medicine. 2011; 31(4): 631–647.

4. Ward E, DeSantis C, Robbins A, Kohler B, Jemal A. Childhood and adolescent cancer statistics. CA: A Cancer Journal for Clinicians. 2014; 64(2): 83–103.

5. Orpha. TAL1TAL bHLH transcription factor 1, erythroid differentiation factor [online] Available at http://www.orpha.net/consor/cgi-bin/Disease_Genes.php?lng=EN&data_id=15576 [Accessed 31 May 2018]

6. Sanda T, Leong WZ. TAL1 as a master oncogene transcription factor in T-cell acute lymphoblastic leukemia. Exp Hematol. 2017; 53: 7–15.

7. Han X, Bueso-Ramos CE. Precursor T-cell acute lymphoblastic leukemia/lymphoblastic lymphoma and acute biphenotypic leukemias. Am J Clin Pathol. 2007; 127(4): 528–44.

8. Fasseu M, Aplan PD, Chopin M, Boissel N, Bories JC, Soulier J, von Boehmer H, Sigaux F, Regnault A. p16INK4A tumor suppressor gene expression and CD3epsilon deficiency but not pre-TCR deficiency inhibit TAL1-linked T-lineage leukemogenesis. Blood. 2007; 110(7): 2610–9.

9. Genecards. TAL1 Gene – Disorders. [online] Genecards.org. Available at: http://www.genecards.org/cgi-bin/carddisp.pl?gene=TAL1 [Accessed 31 May 2018].

10. Baer, R. TAL1, TAL2 and LYL1: a family of basic helix-loop-helix proteins implicated in T cell acute leukaemia. Semin Cancer Biol (1993);4(6):341–7.

11. Correia, N., Melão, A., Póvoa, V., Sarmento, L., de Cedrón, M., Malumbres, M., Enguita, F. and Barata, J. microRNAs regulate TAL1 expression in T-cell acute lymphoblastic leukemia. Oncotarget (2016);7(7):8268–81.

12. Cardoso, B., de Almeida, S., Laranjeira, A., Carmo-Fonseca, M., Yunes, J., Coffer, P. and Barata, J. TAL1/SCL is downregulated upon histone deacetylase inhibition in T-cell acute lymphoblastic leukemia cells. Leukemia (2011);25(10):1578–1586.

13. Brown L, Cheng JT, Chen Q, Siciliano MJ, Crist W, Buchanan G, et al. Site-specific recombination of the tal-1 gene is a common occurrence in human T cell leukemia. EMBO J (1990);9(10):3343–51.

14. Bash RO, Hall S, Timmons CF, Crist WM, Amylon M, Smith RG, et al. Does activation of the TAL1 gene occur in a majority of patients with T-cell acute lymphoblastic leukemia? A pediatric oncology group study. Blood (1995);86(2):666–76.

15. Carroll AJ, Crist WM, Link MP, Amylon MD, Pullen DJ, Ragab AH, et al. The t(1;14)(p34;q11) is nonrandom and restricted to T-cell acute lymphoblastic leukemia: a Pediatric Oncology Group study. Blood (1990);76(6):1220–4.

16. Huret, JL., Labastie, MC., Huret, JL., Labastie MC. TAL1 (T-cell acute leukemia 1). Atlas Genet Cytogenet Oncol Haematol. (1998);2(2):47–48.

17. Collins FS, Brooks LD, Chakravarti A, A DNA polymorphism discovery resource for research on human genetic variation. Genome Res (1998);8:1229–1231.

18. Lander ES. The new genomics: global views of biology. Science. 1996.

19. SNP – NCBI. [online] Available at http://www.ncbi.nlm.nih.gov/snp [Accessed in March 2018]

20. Genemania. [online] Available at http://pages.genemania.org/. [Accessed in March 2018]

21. Nahla E. Abdelraheem, Marwa Mohamed Osman, Osama Muhieldin Elgemaabi, Afra Abdelhamid Fadl Alla, Mosab Mohamed Ismail, Soada Ahmed Osman, Aisha Ismail Ibrahim, Nihad Elsadig Babekir4, Salwa Osman Mekki, Mohamed A. Hassan. Computational Analysis of Deleterious Single Nucleotide Polymorphisms (SNPs) in Human MutS Homolog6 (MSH6) Gene American Journal of Bioinformatics Research (2016);6(2):56–97

22. Ivan Adzhubei, Daniel M. Jordan, Sunyaev S.R. Predicting Functional Effect of Human Missense Mutations Using PolyPhen-2. Current Protocols in Human Genetics. 7(20):1–41.

23. Capriotti E, Fariselli P, Casadio R I-Mutant2.0: predicting stability changes upon mutation from the protein sequence or structure. Nucleic Acids Res (2005);33:W306–10.

24. Capriotti E, Calabrese R and Casadio R. Predicting the insurgence of human genetic diseases associated to single point protein mutations with support vector machines and evolutionary information. Bioinformatics (2006);22: 2729–2734.

25. Capriotti E, Fariselli P, Calabrese R. and Casadio R. Predicting protein stability changes from sequences using support vector machines. Bioinformatics (2005);21(Suppl 2): ii54–ii58

26. Altschul S. F, Madden T. L, Schaffer A. A, Zhang J, Zhang. Z, Miller. W and Lipman D. J. Gapped. BLAST and PSI-BLAST: a new generation of protein database search programs. Nucleic Acids Research (1997);25: 3389–3402.

27. Jesse D, Ziebarth YC, Anindya Bhattacharya and Anlong Chen. PolymiRTS Database 2.0: linking polymorphisms in microRNA target sites with human diseases and complex traits. Nucleic Acids Research (2012);40(Database issue):D216–D221.

28. Venselaar, H., T. A. te Beek, R. K. Kuipers, M. L. Hekkelman and G. Vriend. Protein structure analysis of mutations causing inheritable diseases. An e-Science approach with life scientist friendly interfaces. BMC Bioinformatics (2010);11(1):548.

29. Kelley LA et al. The Phyre2 web portal for protein modeling, prediction and analysis. Nature Protocols. 2015; 10: 845–858.

30. Källberg M, Wang H, Wang S, Peng J, Wang Z, Lu H, Xu H. Temple-based protein structure modelling using the RaptorX website server. Nature Protocols. 2012; 7: 1511–1522.

31. http://www.cgl.ucsf.edu/chimera/. Accessed in March 2018

32. Goddard TD1, Huang CC, Ferrin. TE Software extensions to UCSF chimera for interactive visualization of large molecular assemblies. J Structure (2005); 13(3):473–82.

33. Blom N, Gammeltoft S, Brunak S. Sequence and structure-based prediction of eukaryotic protein phosphorylationsites. J Mol Biol. 1999; 294: 1351–1362. https://doi.org/10.1006/jmbi.1999.3310 PMID:10600390

34. Radivojac P, Vacic V, Haynes C, Cocklin RR, Mohan A, et al. (2010) Identification, analysis, and prediction of protein ubiquitination sites. Proteins 78: 365–380.

35. R. Rajasekaran, C. G. PriyaDoss, C. Sudandiradoss, K. Ramanathan, and R. Sethumadhavan, “In silico analysis of structural and functional consequences in p16INK4A by deleterious nsSNPs associated CDKN2A gene in malignant melanoma,” Biochimie (2008);90(10):1523–1529.

36. R. Rajasekaran and R. Sethumadhavan, “Exploring the cause of drug resistance by the detrimental missense mutations in KIT receptor: computational approach,” Amino Acids (2010):39(3):651–660.

37. R. Rajasekaran, C. Sudandiradoss, C. G. P. Doss, and R. Sethumadhavan, “Identification and in silico analysis of functional SNPs of the BRCA1 gene,” Genomics (2007);90(4):447–452.

38. R. Rajasekaran, C. George Priya Doss, C. Sudandiradoss, K. Ramanathan, R. Purohit, and R. Sethumadhavan, “Effect of deleterious nsSNP on the HER2 receptor based on stability and binding affinity with herceptin: a computational approach,” Comptes Rendus—Biologies (2008);331(6):409–4

39. Alabid T, Kordofani AAY, Atalla B, Altayb HN, Fadla AA, et al. In silico Analysis of Single Nucleotide Polymorphisms (SNPs) in HumanVCAM-1 gene. J Bioinform, Genomics, Proteomics 2016; 1(1): 1004.

40. Alabid T, Kordofani AAY, Atalla B, Altayb HN, Fadla AA, et al. (2016) In silico Analysis of Single Nucleotide Polymorphisms (SNPs) in HumanVCAM-1 gene. JBioinform, Genomics, Proteomics 1(1): 1004.

41. Elshaikh AAF, Ismaiel MM, Osman MM, Shokri SAI et al. Computational Analysis of Single Nucleotide Polymorphism (SNPs) in Human GRM4 Gene. American Journal of Biomedical Research, 2016, Vol. 4, No. 3, 61–73

42. Fadlalla Elshaikh AAE, Elmahdi Ahmed MT, Daf Alla TIM, Mogammed Elbasheer AS, Ahmed AA, et al. (2016) Computational Analysis of Single Nucleotide Polymorphism (Snps) In Human MYC Gene. J Bioinform, Genomics, Proteomics 1(3): 1011.

43. Arshad M, Bhatti A, John P (2018) Identification and in silico analysis of functional SNPs of human TAGAP protein: A comprehensive study. PLoS ONE 13(1): e0188143

44. Hoang, T., Lambert, J. and Martin, R. SCL/TAL1 in Hematopoiesis and Cellular Reprogramming. Current Topics in Developmental Biology (2016); 188:163–204.

45. Porcher, C., Chagraoui, H. and Kristiansen, M. SCL/TAL1: a multifaceted regulator from blood development to disease. Blood (2017); 129(15):2051–2060.

46. Bae, J., Kim, H., Ban, J., Park, H., Kim, S., Kang, S., Park, J., Kim, J. and Chung, J. Association between polymorphisms of TAL1 gene and schizophrenia in a Korean population. Psychiatric Genetics (2012);22(1):50.

47. Dai C, Gu W. p53 post-translational modification: deregulated in tumorigenesis. Trends Mol Med. 2010; 16: 528–536. https://doi.org/10.1016/j.molmed.2010.09.002 PMID: 20932800

48. Shiloh Y, Ziv Y. The ATM protein kinase: regulating the cellular response to genotoxic stress, and more. Nat Rev Mol Cell Biol. 2013; 14: 197–210.

49. Ciesla J, Fraczyk T, Rode W. Phosphorylation of basic amino acid residues in proteins: important but easily missed”. Acta Biochim Pol. 2011; 58: 137–147. PMID: 21623415

50. Glickman MH, Ciechanover A (April 2002). “The ubiquitin-proteasome proteolytic pathway: destruction for the sake of construction”. Physiological Reviews. 82 (2): 373–428. doi:10.1152/physrev.00027.2001. PMID 11917093.

51. Mukhopadhyay D, Riezman H (January 2007). “Proteasome-independent functions of ubiquitin in endocytosis and signaling”. Science. 315 (5809): 201–5. doi:10.1126/science.1127085. PMID 17218518.

52. Schnell JD, Hicke L (September 2003). “Non-traditional functions of ubiquitin and ubiquitin-binding proteins”. The Journal of Biological Chemistry. 278 (38): 35857–60. doi:10.1074/jbc.R300018200. PMID 12860974.

